# Habitat fragmentation enhances microbial collective defence

**DOI:** 10.1101/2024.03.20.585867

**Authors:** Nia Verdon, Ofelia Popescu, Simon Titmuss, Rosalind J. Allen

## Abstract

Microbes often inhabit complex, spatially partitioned geometries such as host tissue or soil, but the effects of habitat fragmentation on microbial infection dynamics and ecology are poorly understood. Here we investigate how habitat fragmentation impacts a prevalent microbial collective defence mechanism: enzymatic degradation of an environmental toxin. Using a theoretical model, we predict that habitat fragmentation can strongly enhance the collective benefits of enzymatic toxin degradation. For the clinically relevant case where *β*-lactamase producing bacteria mount a collective defence by degrading a *β*-lactam antibiotic, we find that realistic levels of habitat fragmentation can allow a population to survive antibiotic doses that would far exceed those required to kill a non-fragmented population. This “habitat-fragmentation rescue” is a stochastic effect that originates from variation in bacterial density among different subpopulations and demographic noise. In contrast, the stochastic effects of habitat fragmentation are weaker in a model of collective enzymatic nutrient foraging. Our model suggests that treatment of a spatially complex, fragmented infection showing collective resistance may be far less effective than expected based on bulk population assumptions. This may help to explain lack of correlation between lab-measured antibiotic susceptibility values and clinical treatment success.

## Introduction

Microbial communities commonly inhabit spatially fragmented habitats, such as particles or aggregates in aqueous environments (***Blackburn et al., 1998; Dann et al., 2014***), plant roots (***Edwards et al., 2015***) or leaf surfaces (***Monier and Lindow, 2004; Grinberg et al., 2019***), crypts in the gut lining (***Welch et al., 2017***), pores in the soil (***Bickel and Or, 2020; Raynaud and Nunan, 2014***), in the skin (***Conwill et al., 2022***) or, for intracellular pathogens, the interior of host cells (***Ray et al., 2009***). Even in the absence of physical habitat fragmentation, slow diffusion of nutrients and/or signals can lead to effective fragmentation (***Dal Co et al., 2020***). Habitat fragmentation can significantly affect microbial interactions (***Boedicker et al., 2009; Connell et al., 2014; Geyrhofer and Brenner, 2020; Batsch et al., 2024***), community assembly (***Hansen et al., 2016; Hsu et al., 2019***) and population dynamics (***Wu et al., 2022; Mant et al., 2024***), but mechanistic understanding of these effects is lacking. Here we investigate the mechanisms by which habitat fragmentation alters microbial interactions, by developing a mathematical model in which a microbial population is systematically partitioned into smaller sub-habitats, while keeping the total population size and volume the same (Fig. 1). Using this approach, we show how habitat fragmentation can strongly affect a prevalent and clinically class of microbial behaviour: collective defence by enzymatic degradation of an environmental toxin.

**Figure 1.**
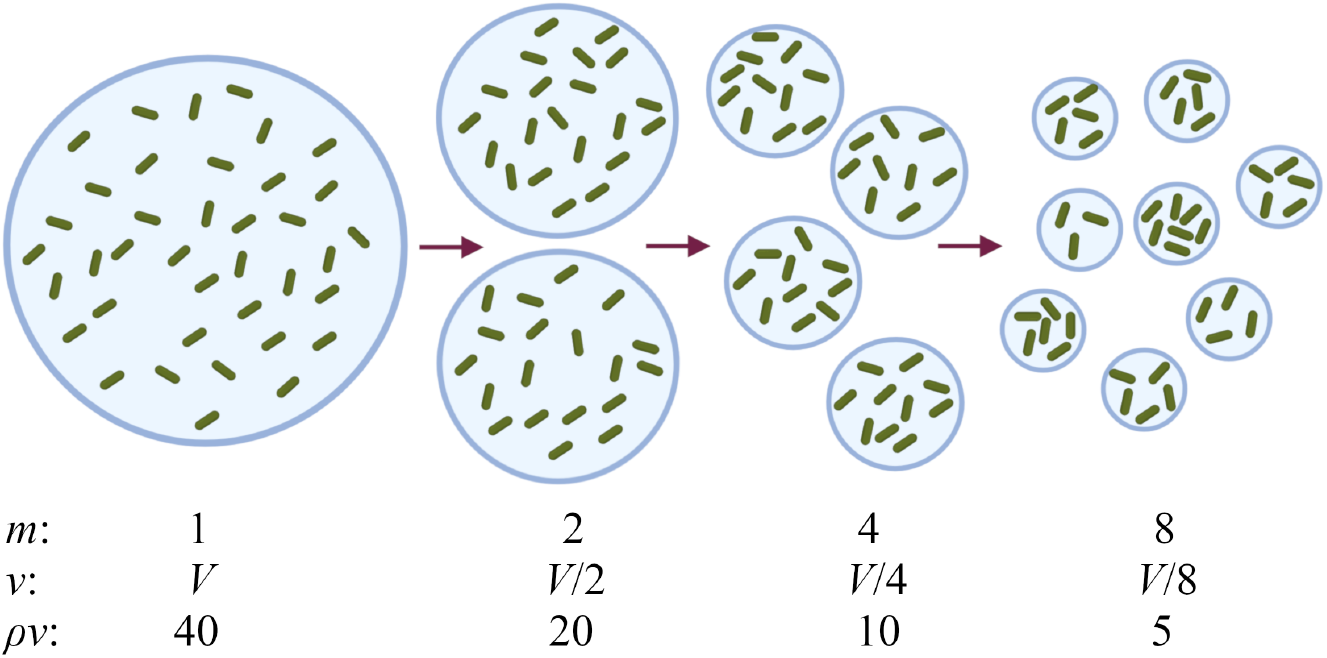
Modelling habitat fragmentation by systematically partitioning a microbial population. A population of microbes of density *ρ*, contained in volume *V*, is partitioned into smaller subvolumes of volume *v* = *V* /*m*. For degree of partitioning *m*, the average number of microbes per subvolume is *ρv*. Microbes are distributed stochastically among the subvolumes, so that the number of microbes in a given subvolume follows a Poisson distribution. As an exemplar, in the diagram we use *ρv* = 40 for *m*=1.

Many microbes produce enzymes that modify their local environment; examples include antibiotic-degrading enzymes such as *β*-lactamases (***Tooke et al., 2019***), cellulases that allow digestion of plant matter (***Thapa et al., 2020***), hydrolases that degrade marine polysaccharides (***Arnosti, 2011***), sucrose digestion by yeast invertases (***Gore et al., 2009***), and the production of virulence factors such as elastase that cause tissue damage (***Galloway, 1991***). Enzymatic modification of the environment is a collective behaviour, since the effects are shared by nearby microbes. Indeed, enzyme production has been analysed in the framework of social evolution theory, which addresses how enzyme production can be maintained evolutionarily in the face of cheater non-producers who reap the benefits but do not pay the cost of enzyme production (***Vega and Gore, 2014; Yurtsev et al., 2013; Mizrahi et al., 2022***). For mixed populations of cooperators and cheaters, habitat fragmentation can favour the evolution of cooperators through kin selection, although it may also increase the level of local competition between cooperators (***Tekwa et al., 2017***).

Here, we consider a clonal population, composed entirely of enzyme producers (i.e. cooperators). We focus on the production of *β*-lactamase enzymes by antibiotic-resistant bacteria (***Tooke et al., 2019***). These enzymes degrade *β*-lactams, which are among the most commonly prescribed classes of antibiotic (***Bush and Bradford, 2016; Tooke et al., 2019***). Therefore, *β*-lactamase producing antibiotic-resistant infections present a severe clinical problem, and improved strategies to combat them could have significant impact (***Tooke et al., 2019***). Consistent with the concept of *β*-lactamase production as a collective defence strategy against the antibiotic, *β*-lactamase producing bacteria often show an inoculum effect, whereby populations with high initial bacterial density survive, while those with low initial density are killed (***Vega and Gore, 2014; Brook, 1989; Meredith et al., 2015***). Recent mathematical modelling has shown that this effect can be understood as a race for survival, which depends on the relative timescales for antibiotic killing versus antibiotic degradation (***Geyrhofer et al., 2023***). However, the role of habitat fragmentation has not yet been considered.

Here, we present a theoretical modelling approach which we use to predict the effects of habitat fragmentation on microbial enzyme-producers. Our main result is that, for microbes that engage in collective defence via enzymatic degradation of a toxin (such as *β*-lactam degradation), habitat fragmentation can dramatically increase the probability of survival. This suggests that antibiotic treatment of a fragmented *β*-lactamase producing infection may be far less effective than would be expected based on the measured susceptibility (i.e. the minimum inhibitory concentration, or MIC value). In other words, habitat fragmentation can rescue an infection, allowing it to survive and regrow after the antibiotic is degraded. In contrast, we find that the effects of fragmentation are weaker for a microbial population that uses enzymes to boost growth by releasing nutrient from the environment. The effects predicted here are intrinsically stochastic, arising from the variability in initial population densities among the partitioned subpopulations, as well as from demographic stochasticity in microbial growth and killing dynamics.

Enzymatic modification of the environment by microbes is ubiquitous, from clinical infections to biotechnology and biogeochemistry (***Tooke et al., 2019; Thapa et al., 2020; Arnosti, 2011; Gore et al., 2009; Galloway, 1991; Zhang et al., 2021; Hermenau et al., 2020***). Our study shows how the intrinsic stochasticity associated with habitat fragmentation can alter the ecology of these interactions.

## Results

### Collective defence via enzymatic degradation

We first consider a microbial population that is exposed to an environmental toxin – specifically, a population of *β*-lactamase producing bacteria exposed to a *β*-lactam antibiotic. As a simple model, we suppose that the population (contained in volume *V*) grows exponentially if the antibiotic concentration is below a threshold *a*_th_ but dies exponentially if the antibiotic concentration is above the threshold. The threshold concentration *a*_th_ corresponds to the single-cell minimum inhibitory concentration (scMIC) (***Artemova et al., 2015***). In addition, the microbes produce antibiotic-degrading *β*-lactamase enzyme: this causes the antibiotic concentration to decrease with time. The microbial population size *N*(*t*) is described by the following dynamical equation:

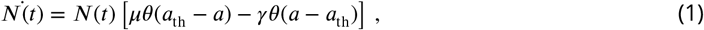

where *θ*(*x*) denotes the Heaviside step function (*θ*(*x*) = 1 if *x* ≥ 0 and zero otherwise), *μ* is the growth rate for low antibiotic *a*(*t*) < *a*_th_, and *γ* is the death rate for high antibiotic *a*(*t*) > *a*_th_. The antibiotic is degraded according to:

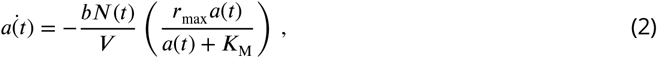

where *b* is the number of enzyme molecules per cell (i.e. the enzyme concentration is *bN*/*V*) and each enzyme degrades antibiotic according to Michaelis-Menten kinetics with parameters *r*_max_ and *K*_M_ (note that if the enzyme remains partially or wholly within the bacterial cell, the *r*_max_ parameter also implicitly accounts for antibiotic transport across the cell boundary (***Dugatkin et al., 2005; Geyrhofer et al., 2023***)). We assume that the initial antibiotic concentration, *a*_init_, is high, *a*_init_ > *a*_th_. Similar models have been proposed in previous work (***Yurtsev et al., 2013; Mizrahi et al., 2022; Geyrhofer et al., 2023***); more detailed models of antibiotic inhibition lead to qualitatively equivalent results (***Geyrhofer et al., 2023***).

In our model, two distinct outcomes are possible (Fig. 2a,b). The microbial population may be killed outright, or it may reduce the antibiotic concentration below the threshold *a*_th_, and then regrow. This phenomenon has previously been described as a race for survival, since the outcome depends on the relative timescales of killing and antibiotic degradation (***Geyrhofer et al., 2023***); Fig. 2c. The population will survive and regrow if its initial density is greater than a critical value *ρ*^∗^ that depends on the initial antibiotic concentration (***Geyrhofer et al., 2023***):

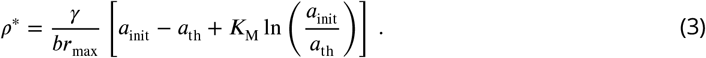

(for derivation of Eq. 3, see Supplementary Theoretical Derivations, Eqs. S6-S14). The line *ρ*^∗^(*a*_init_) defined by Eq. 3 defines an ecological phase boundary that separates regions of parameter space where the microbial population is killed from regions where it survives and regrows (Fig. 2d).

**Figure 2.**
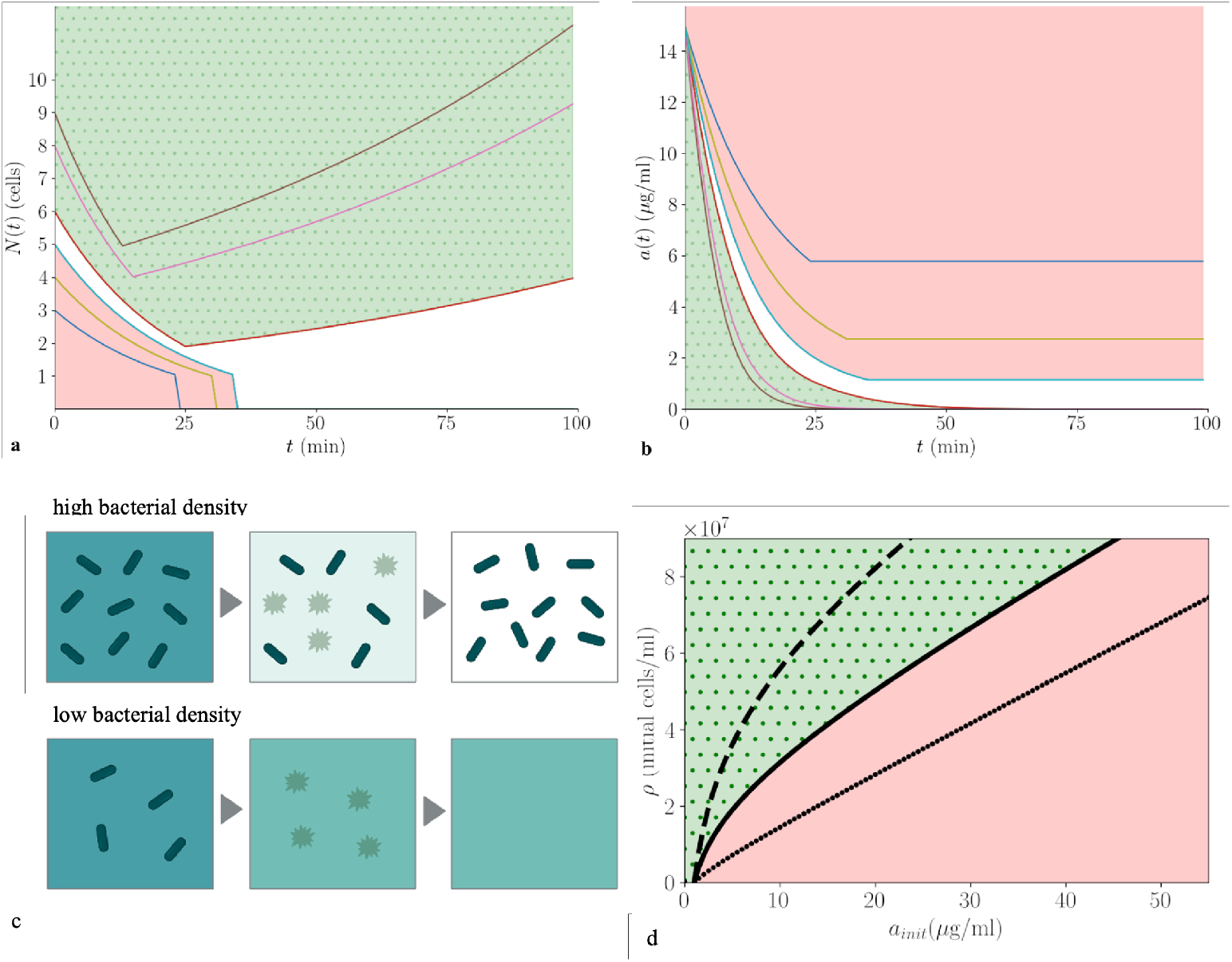
Collective enzymatic defence: population density threshold for survival. (a,b): Predictions of the model, Eqs. 1 and 2, for the microbial population size *N*(*t*) and the antibiotic concentration *a*(*t*) (panels (a) and (b) respectively), for a range of values of the initial microbial population size *N*_init_ (coloured lines; *N*_init_ ranges between 3 and 9). The colour coding is consistent between (a) and (b). The red shaded region indicates death of the microbial population; the green shaded region indicates survival and regrowth of the population. The parameters are as follows: *br*_max_ = 3.5 × 10^−8^ *μ*g/cell/min (***Yurtsev et al., 2013***), *K*_M_ = 6.7 *μ*g/ml (***Yurtsev et al., 2013***), *a*_th_ = 1 *μ*g/ml (***Yurtsev et al., 2013***), *a*_init_ = 15 *μ*g/ml, *μ* = 0.01 /min (***Taylor et al., 2022***), *γ* = 0.045 /min (***Yurtsev et al., 2013***), *V* = 100 pl (***Taylor et al., 2022***). (c): Schematic showing the two ecological outcomes. The background colour indicates antibiotic concentration (dark = high, light = low) while the dark green objects represent bacteria, which can be alive (rod-shaped) or dead (faint stars). At high bacterial density, antibiotic is degraded below the threshold *a*_th_ = 1 before the population is entirely killed, therefore the population recovers; at low bacterial density the entire population is killed before the antibiotic concentration reaches the threshold. (d): Phase diagram showing the predicted outcome (killing or survival and regrowth; red and green, respectively) as a function of initial bacterial density *ρ* and initial antibiotic concentration *a*_init_. The lines indicate the critical density *ρ*^∗^, Eq. 3, for different values of the Michaelis-Menten parameter *K*_M_: solid line *K*_M_ = 6.7 *μ*g/ml (***Yurtsev et al., 2013***), dashed line *K*_M_ = 15 *μ*g/ml, dotted line *K*_M_ = 1 *μ*g/ml.

The shape of the ecological phase boundary reveals an inoculum effect: populations with a higher initial density can survive at higher initial antibiotic concentrations (***Brook, 1989***) (Fig. 2d).

This arises from the collective nature of antibiotic degradation (***Vega and Gore, 2014; Yurtsev et al., 2013; Mizrahi et al., 2022; Geyrhofer et al., 2023***). The qualitative nature of the inoculum effect depends on the kinetic parameters of the enzyme. If *K*_*M*_ is small (*a*_init_ ≫ *K*_*M*_), the phase boundary is linear (Fig. 2d, dots), but if *K*_*M*_ is large (*a*_init_ ≪ *K*_*M*_), it becomes logarithmic (Fig. 2d, dashes; Eq. 3).

### In a fragmented habitat, subpopulation killing is stochastic

We now ask how habitat fragmentation (e.g. within skin pores, or crypts in the gut lining) affects the fate of a microbial population engaged in collective defence, under lethal conditions where a well-mixed population would be killed. Using as our example a population of *β*-lactamase producing bacteria exposed to an antibiotic concentration well above the MIC, we now suppose that the total volume *V*, containing a population of density *ρ*, is partitioned into *m* subvolumes of volume *v* = *V* /*m* (***Wu et al., 2022***) (Fig. 1). The average (initial) number of microbes per subvolume is then 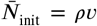. However, the subvolumes are filled stochastically, so the number of microbes *N*_*i*,init_ in subvolume *i* is sampled from a Poisson distribution with probability 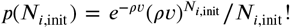. Poisson statistics have indeed been observed for encapsulation of bacteria in microfluidic droplets (***Collins et al., 2015; Barizien et al., 2019; Taylor et al., 2022***). Each subvolume also contains antibiotic at uniform concentration *a*_init_. Neither microbes nor antibiotic can be exchanged between subvolumes, so that the microbe-antibiotic dynamics evolve independently in each subvolume. Because different subvolumes have stochastically different initial subpopulation sizes, we anticipate different outcomes among replicate subvolumes (Fig. 3a). We model the dynamics in each sub-volume deterministically via Eqs. 1 and 2 (although later we also consider stochastic birth-death dynamics).

**Figure 3.**
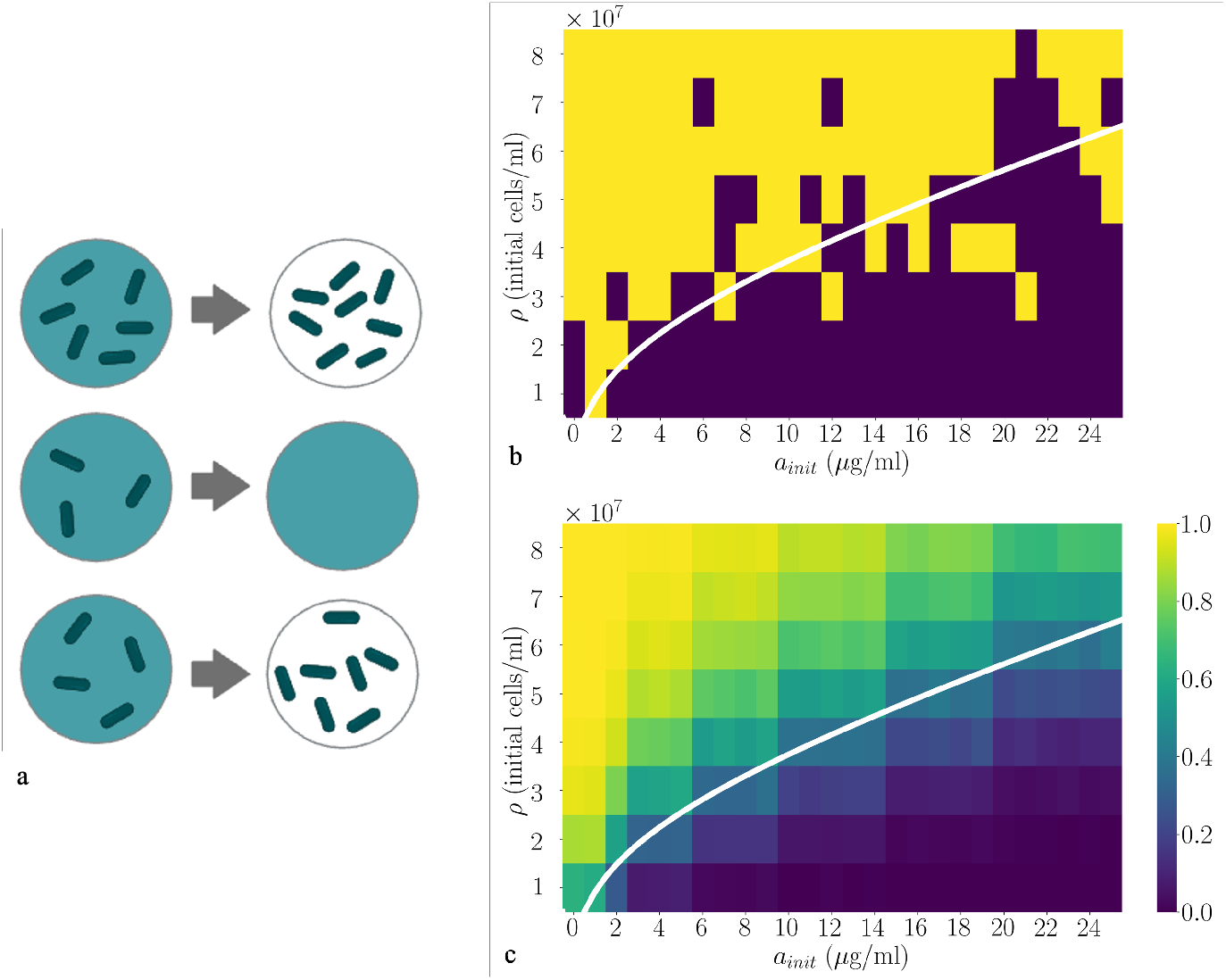
Subpopulation killing is stochastic. (a): Illustration that different replicate subpopulations experience different ecological outcomes depending on their stochastic initial population density. The background colour represents antibiotic concentration (dark = high, light = low) while the dark green objects represent live bacteria. A subpopulation that is below the threshold density for survival *ρ*^∗^ is killed, while subpopulations with higher initial density survive and grow. (b): The model of Eqs. 1 and 2 is simulated for a single subpopulation with Poisson distributed initial population size, for a range of values of the initial antibiotic concentration *a*_init_ and the mean population density *ρ*, with *v* = 100 pl. The other parameters are *br*_max_ = 3.5 × 10^−8^*μ*g/cell/min, *K*_M_ = 6.7 *μ*g/ml, *a*_th_ = 1*μ*g/ml, *μ* = 0.01/min, *γ* = 0.045/min (as in Fig. 2). Simulation runs where the subpopulation is killed are shown as dark squares; those where the subpopulation survives and regrows are shown as light squares. (c): Equivalent simulations to those of (b), averaged over 1000subpopulations for each condition. Here the probability *p*_*s*_ of subpopulation survival is indicated by the colour scale. In both (b) and (c) the ecological phase boundary corresponding to *ρ*^∗^(*a*_init_) from Fig. 2d is shown as a white line. For a detailed description of the simulation methods, see Supplementary Numerical Simulation Methods.

The relevant ecological outcome here is the survival vs killing of the fragmented microbial population – in antibiotic treatment, even a few surviving microbes can regrow and cause disease recurrence. We therefore calculate the probability that the entire microbial population is killed, or its complement, the “survival probability”, which is the probability that any microbes within the population survive the treatment.

We first consider the subpopulation of microbes contained within a single subvolume *v*. In our model, the fate of the subpopulation is determined by its initial density (Eq. 3 and Fig. 2d); it survives if *N*_*i*,init_ /*v* > *ρ*^∗^. The probability *p*_*s*_ for this condition is controlled by the Poisson distribution of initial subpopulation sizes: 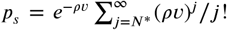, where *N*^∗^ = ⌈*ρ*^∗^*v*⌉. Indeed, numerical simulations of the model dynamics for a single subpopulation with Poisson distributed initial population size show stochastic outcomes (Fig. 3a,b). For antibiotic concentrations and (average) microbial densities below the ecological phase boundary, for which the bulk, non-fragmented population is killed (Fig. 2d), some of the stochastically simulated subpopulations survive. Conversely, above the phase boundary, where a bulk population survives, some stochastically simulated subpopulations are killed (Fig. 3b). Averaging these results over many stochastic simulation runs (Fig. 3c) highlights that there is a non-zero probability of survival in regions of parameter space that are below the phase boundary and, conversely, a non-zero probability of killing in regions above the phase boundary. Therefore, stochasticity in the initial population density can change the ecological outcome for individual subpopulations.

### Habitat fragmentation can rescue a microbial population in a lethal environment

We now consider the fate of the entire microbial population, consisting of *m* independent subpopulations. The probability that the entire population is killed is given by

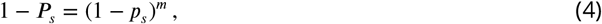

where *P*_*s*_ is the total population survival probability, i.e. the probability that *any* of the subpopulations survives and regrows.

Habitat fragmentation strongly increases the probability of population survival. For conditions below the ecological phase boundary, for which a bulk population would be killed, simply fragmenting the population into subvolumes (i.e. increasing *m* or equivalently decreasing *ρv* without changing the total population size or the total volume) can ensure its survival (Fig. 4a). Thus the model predicts that a fragmented infection can survive antibiotic treatment doses that would be sufficient to kill a non-fragmented infection. To explore this point more explicitly, we model a *β*-lactamase producing bacterial infection consisting of 5000 cells at density 5 × 10^7^ cells/ml in a volume 10^8^*μ*m^3^ (Fig. 4b). In an unfragmented habitat this infection will be killed at antibiotic concentrations above the single-cell MIC (in this model, 1 *μ*g/mL). However if the habitat is fragmented into 100 sub-habitats of volume 10^6^*μ*m^3^ (consistent with lung alveolae (***Ochs et al., 2004***) or hair follicles (***Blume-Peytavi et al., 2008***)), with on average 50 cells per subhabitat, the model predicts that even an antibiotic concentration of 35 *μ*g/mL is insufficient to eliminate the infection (Fig. 4a, arrow and Fig. 4b).

**Figure 4.**
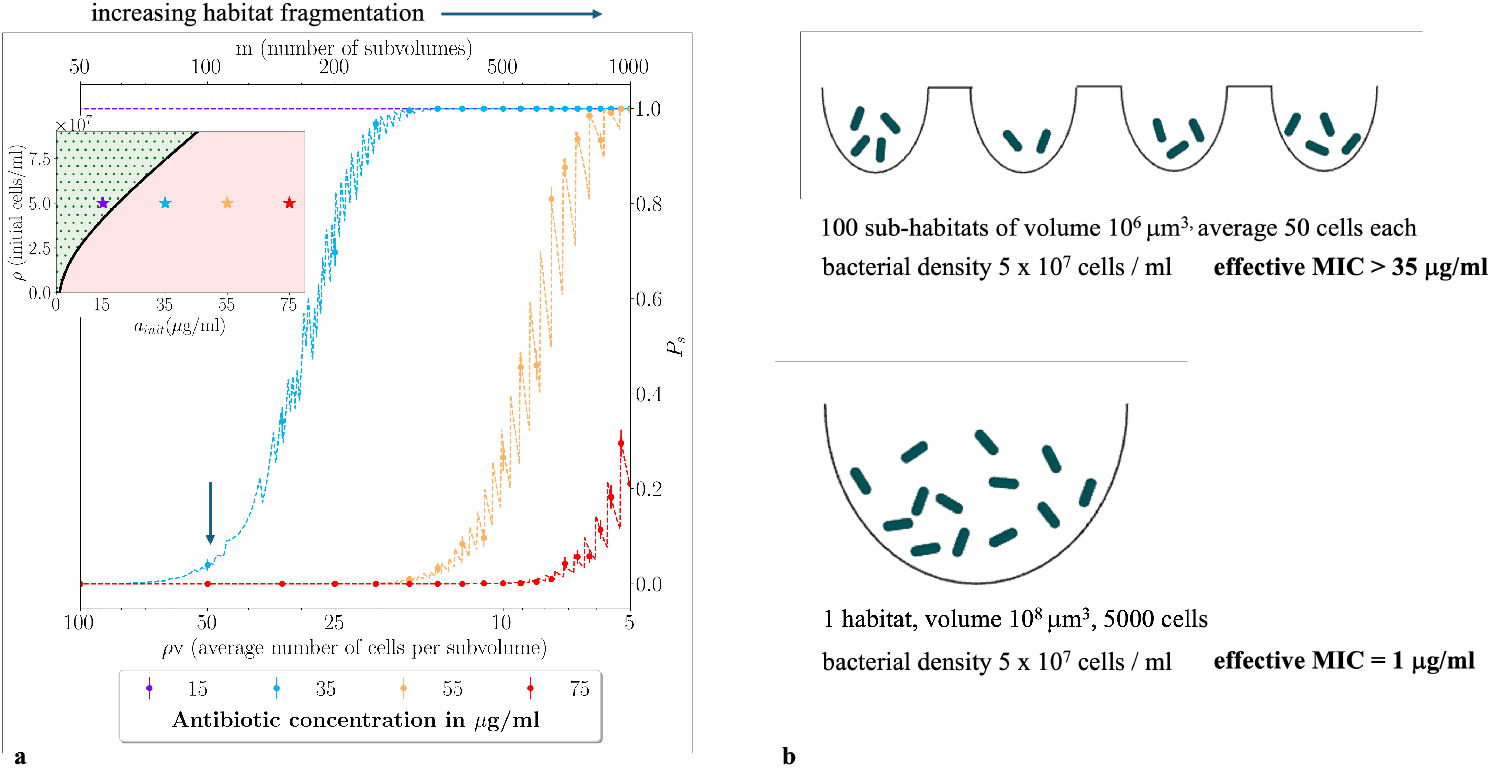
Habitat fragmentation can rescue a microbial population in a lethal environment. (a): Total population survival probability *P*_*s*_ increases from zero to 1 as the habitat becomes more fragmented, for lethal parameter combinations for which a bulk population would be killed. *P*_*s*_ is plotted as a function of *m* (top axis), or the average subpopulation size *ρv* (bottom axis), for a range of initial antibiotic concentrations *a*_init_ (shown by colour) and mean population density *ρ* = 5 × 10^7^ cells/ml. The data symbols show results of numerical simulations of the model of Eqs. 1 and 2 with a Poisson distribution of initial subpopulation sizes (each data point is the mean of 1000 simulation runs; error bars show 95% confidence interval, assuming a binomial distribution), while the dashed lines show theoretical predictions obtained from Eq. 4. For the lowest antibiotic concentration (15 *μ*g/ml) the bacterial population always survives. For higher concentrations (35 *μ*g/ml, 55 *μ*g/ml, 75 *μ*g/ml), the population is killed for low habitat fragmentation but starts to survive as the degree of habitat fragmentation increases. The arrow indicates the condition analyzed in panel (b). The zig-zagging in the theoretical lines arises because of the discreteness of individual microbes, reflected in the bound *N*^∗^ = ⌈*ρ*^∗^*v*⌉ in the sum that features in the expression for *p*_*s*_. The other parameters are *br*_max_ = 3.5 × 10^−8^*μ*g/cell/min, *K*_M_ = 6.7 *μ*g/ml, *a*_th_ = 1*μ*g/ml, *μ* = 0.01/min and *γ* = 0.045/min (as in Fig. 2). The colour coded points in the inset show the location in the (*a*_init_, *ρ*) phase diagram for each antibiotic concentration, relative to the survival phase boundary; a bulk population survives for 15*μ*g/mL but dies for 35, 55 or 75 *μ*g/mL (*ρ* ∗ values are 4.1 × 10^7^, 7.4 × 10^7^, 1.04 × 10^8^, and 1.3 × 10^8^ cells/ml, respectively). For a detailed description of the simulation approach, see Supplementary numerical simulation methods. (b): Consequences for treatment of fragmented vs non-fragmented infections. Using the same parameters as in (a), we compare a bacterial population of density 5 × 10^7^ cells/ml that is either fragmented into ∼ 100 sub-habitats each of volume 10^6^*μ*m^3^ (top; indicated by the arrow in (a)) or contained in an unfragmented habitat of volume ∼ 10^8^*μ*m^3^ (bottom). The unfragmented population is predicted to have an effective MIC equal to the single cell MIC (here 1 *μ*g/mL), while the fragmented population has a much higher effective MIC of 35 *μ*g/mL.

In our model, this “fragmentation rescue” phenomenon arises from the stochastic distribution of microbes among the subvolumes. Because the microbes are distributed stochastically, some subvolumes have an initial population density higher than the survival threshold *ρ*^∗^ (i.e. above the phase boundary) even though the average density *ρ* is below the phase boundary. For a Poisson distribution of cell numbers, the coefficient of variation of the population density increases with the degree of partitioning *m*^1^; hence the chance of survival is increased by habitat fragmentation. Three distinct antibiotic killing regimes can be distinguished, for different degrees of habitat fragmentation. For low fragmentation (small *m* or large *ρv* in Fig. 4a), stochasticity is not important and the infection is killed (*P*_*s*_ ≈ 0). As the degree of fragmentation increases, stochasticity becomes relevant and there is a significant probability of survival; here the model predicts an approximately linear relation between log *P*_*s*_ and the subvolume size *v* (Supplementary Text and Fig. S1). In the extreme fragmentation regime (very large *m* or very small *ρv*), the model predicts that the entire population survives, even for very high antibiotic concentration (Fig. 4a; see also Supplementary Fig. S3a). In this regime, the subvolume size becomes small enough that the cell density exceeds the critical value *ρ*^∗^ even for a single microbe (1/*v* > *ρ*^∗^). However, our model is likely to be unrealistic in this limit (see Supplementary text and Fig. S3 for further discussion).

For bacterial populations that do survive and regrow, spatial partitioning also influences the eventual population size, since subpopulations with stochastically higher initial population size start to regrow earlier (Supplementary text and Fig. S2a).

### Other sources of stochasticity can also lead to habitat-fragmentation rescue

So far, the only source of stochasticity in our analysis has been the random partitioning of microbes between subpopulations, since we used a deterministic model, Eqs. 1 and 2, for microbial growth and death. However, other sources of stochasticity may be significant in small populations, including demographic stochasticity arising from the randomness of microbial birth and death events (***Allen and Waclaw, 2019; Khatri et al., 2012***), and phenotypic or genotypic heterogeneity between individual microbial cells (***Ackermann, 2015***).

We explored the role of demographic stochasticity by including stochastic birth and death processes in our model (using a modified version of the Gillespie algorithm (***Gillespie, 1976***); see Supplementary text). Our simulations show that intrinsic stochasticity in the microbial killing dynamics can also rescue a population in a fragmented habitat (Fig. S2) (***Teimouri and Kolomeisky, 2019***). For the parameter set chosen here (representative of collective *β*-lactamase degradation in a microfluidic droplet setup (***Taylor et al., 2022***)), the contribution of demographic stochasticity is approximately equal in magnitude to that of stochastic partitioning, with maximal effect when both sources of stochasticity are present (Figs. S2 and S4).

### Habitat fragmentation inhibits growth in a non-lethal environment

Interestingly, the effects of habitat fragmentation are qualitatively different when the environment is non-lethal, i.e. for low antibiotic concentrations or high cell densities, above the phase boundary for survival, *ρ* > *ρ*^∗^ (green region of Fig. 2d). In this case, habitat fragmentation is actually detrimental to the bacterial population. Our simulations show that the subpopulation survival probability *p*_*s*_ decreases with the degree of fragmentation for low antibiotic concentrations (Fig. 5a). This is because even if the average population density is above the survival threshold *ρ*^∗^, some subpopulations have initial population density below the threshold and are killed by the antibiotic. These subpopulations will not contribute to population growth (Fig. 5b).

**Figure 5.**
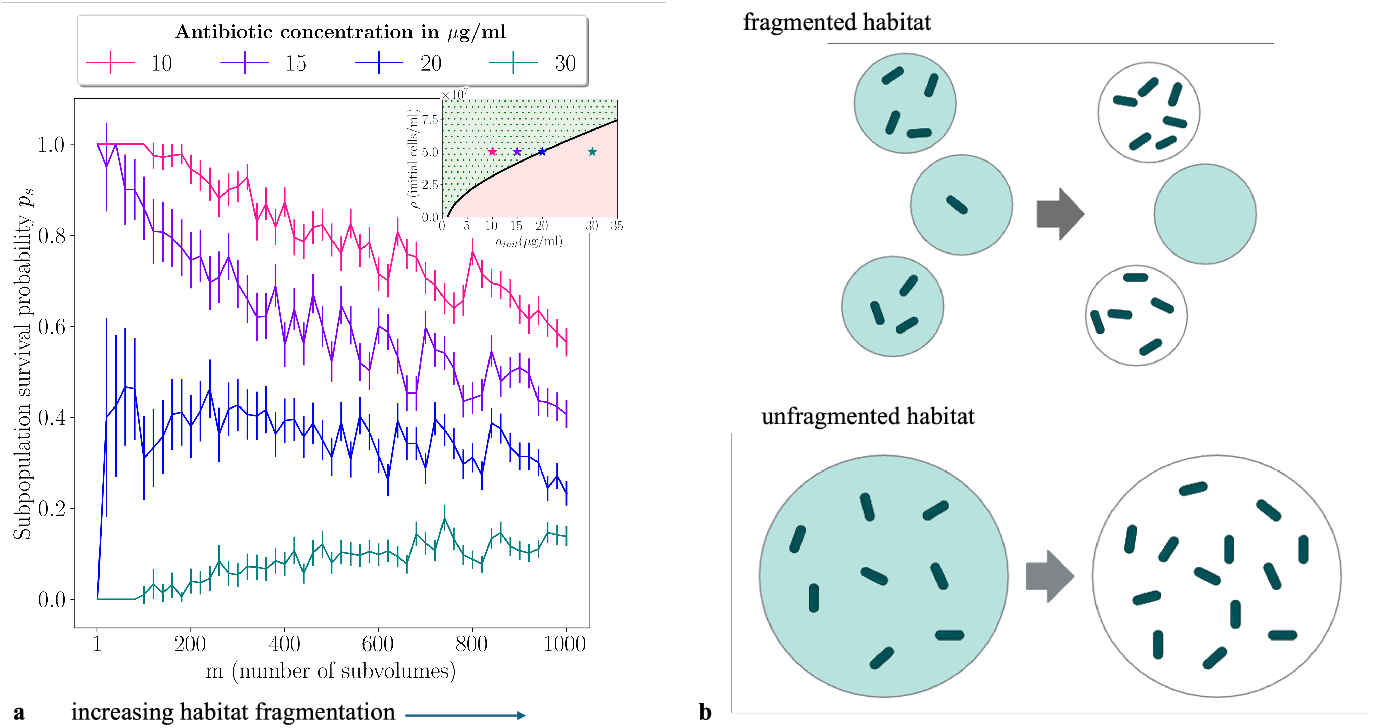
Habitat fragmentation is detrimental for microbes in a non-lethal environment. (a): The probability *p*_*s*_ of survival for any given subpopulation, conditional on the subvolume being initially occupied, is plotted as a function of the degree of habitat fragmentation *m*. The data is obtained from numerical simulations of the model of Eqs. 1 and 2 with a Poisson distribution of initial subpopulation sizes. The mean population density *ρ* = 5 × 10^7^ cells/ml and the other parameters are *br*_max_ = 3.5 × 10^−8^*μ*g/cell/min, *K*_M_ = 6.7 *μ*g/ml, *a*_th_ = 1 *μ*g/ml, *μ* = 0.01/min and *γ* = 0.045 /min (as in Fig. 2). The subpopulation survival probability decreases with habitat fragmentation for low initial antibiotic concentrations, i.e. for non-lethal conditions under which an unfragmented population would survive. The colour coded points in the inset show the location in the (*a*_init_, *ρ*) phase diagram for each antibiotic concentration, relative to the survival phase boundary. Antibiotic concentrations 10 *μ*g/ml and 15 *μ*g/ml are to the left of the phase boundary (non-lethal), 30 *μ*g/ml is to the right of the boundary (lethal) and 20 *μ*g/ml lies right on the boundary. (b): Illustration of how stochastic partitioning of microbes between sub-habitats decreases overall growth under (on average) non-lethal conditions. Sub-populations with low initial density are killed and do not contribute to population growth, even though the conditions are such that an unfragmented population survives.

### Habitat fragmentation can decrease lag times for collective resource foraging via enzymatic degradation

Microbes can modify their environment enzymatically in diverse ways, not limited to detoxification. As a contrasting example, we also modelled a microbial population which uses enzymes to release collective nutritional benefits from its environment. Examples of such behaviour include the breakdown of sucrose by the enzyme invertase produced by yeasts (***Gore et al., 2009***), particulate organic matter breakdown by marine bacteria (***Ebrahimi et al., 2019***) and the breakdown of plant cellulose by microbial cellulase enzymes (***Thapa et al., 2020***). We reasoned that collective resource foraging interactions might also be affected by habitat fragmentation.

To probe this idea, we modified the model of Eqs. 1,2 to represent a microbial population which produces enzymes that release nutrients from the environment, facilitating growth. We suppose that the population cannot grow until the nutrient concentration exceeds a threshold, but once the nutrient threshold is reached it grows exponentially (see Supplementary results and Fig. S5). For an unfragmented population, this model shows an initial lag time while the nutrient concentration is below the threshold, followed by exponential growth once the nutrient reaches the threshold.

Repeating our simulations with systematically increasing degrees of habitat fragmentation, we observed that fragmentation decreases the lag time before population growth starts (Fig. S5). This happens because subpopulations that stochastically have a higher initial population density release environmental nutrients faster, allowing them to initiate growth earlier. In this case the ecological outcome – lag followed by growth – is the same whether or not the habitat is fragmented. Nevertheless, habitat-fragmentation induced lag time decrease could be relevant, for example in the case of a temporally fluctuating environment.

## Discussion

Microbes commonly inhabit fragmented habitats, yet most laboratory studies use well-mixed, unfragmented populations. There is emerging interest in how habitat fragmentation influences microbial interactions, population dynamics and community ecology (***Boedicker et al., 2009; Connell et al., 2014; Geyrhofer and Brenner, 2020; Hansen et al., 2016; Hsu et al., 2019; Wu et al., 2022; Mant et al., 2024; Batsch et al., 2024***), but the overall picture is not yet clear. Here, we show using a theoretical model that enzymatically-mediated collective defence against a chemical toxin can be far more effective in a fragmented habitat than in a well-mixed environment. This is especially relevant in the context of the degradation of clinically relevant *β*-lactam antibiotics by *β*-lactamase enzymes, which are of global importance since they are the primary resistance mechanism against these antibiotics (***Tooke et al., 2019***). Indeed, the World Health Organisation (WHO) and US Center for Disease Control and Prevention (CDC) have identified carbapenemase-producing strains of *Pseudomonas aeruginosa, Acinetobacter baumannii* and Enterobacterales as critical threats (***Tacconelli, 2017; CDC, 2019***). Our work suggests that antibiotic therapy may be even less effective against *β*-lactamase producing strains than suggested by their (already high) MIC values, since in a fragmented environment (such as some human tissues) even doses well above the MIC may fail to eliminate an infection. In our model, this phenomenon originates from the pockets of high local microbial density that can arise in a fragmented environment, even when the average density is much lower. Within such high density pockets, enzyme-producing microbes are protected by their ability to rapidly degrade the antibiotic. Local pockets of high microbial density are indeed observed in soil (***Kuzyakov and Blagodatskaya, 2015***), in aquatic systems (***Blackburn et al., 1998***) and in biofilm infections (***Bjarnsholt et al., 2011***), suggesting that habitat-fragmentation rescue might be a widespread phenomenon.

In our simulations, the distribution of microbial cells among subvolumes was the main source of stochasticity leading to habitat-fragmentation rescue. We also showed that demographic noise in birth/death dynamics can provide an alternative and (for our parameter set) similarly effective source of stochasticity. Our results support the emerging idea that habitat fragmentation increases the variability of ecological outcomes in microbial communities (***Batsch et al., 2024***), and also strengthens the case that stochasticity can play a key role in determining the efficacy of antibiotic treatment (***Balaban et al., 2004; Coates et al., 2018; Deris et al., 2013; Alexander and MacLean, 2020***). Another important source of stochasticity in microbial communities is phenotypic and genotypic heterogeneity among individual microbial cells (***Ackermann, 2015***). Such heterogeneity has been implicated in the dynamics of intracellular bacterial infections (***Avraham et al., 2015; Ortega et al., 2019***); it would be interesting in future to extend our work to include heterogeneity among bacterial cells.

Collective benefits among microbes are often viewed in the context of social evolution theory, where key questions concern the establishment and maintenance of cooperators (here enzyme producers) in the presence of non-producing cheaters who do not pay the fitness cost of production (***West et al., 2006***). In social evolution theory, fragmentation of a population increases the chance for genetically related organisms to share the benefits of cooperation (kin selection) but can also increase competition among related organisms (***Peña and Nöldeke, 2018***). Fragmenting a population into groups of variable size can maintain cooperators via Simpson’s paradox (***Chuang et al., 2009***). Our study shows that, even for a clonal population of cooperators, habitat fragmentation can strongly change the value of a collective benefit, especially in the case of toxin removal. Future work should explore the implications of this insight for mixed populations of cooperators and cheaters.

Our study is based on a simple theoretical model, although more realistic models are expected to show similar behaviour (***Geyrhofer et al., 2023***). Microfluidic technology increasingly allows controlled study of the dynamics of compartmentalised small microbial populations (***Connell et al., 2014; Boedicker et al., 2009; Dal Co et al., 2020; Barizien et al., 2019; Taylor et al., 2022; Park et al., 2011; Batsch et al., 2024***), including their response to antibiotics (***Scheler et al., 2020***). Our study provides a baseline prediction; the reality may be more complex due to phenomena such as morphological or gene regulatory changes in response to antibiotic (***Zhao et al., 2023***) and/or de novo evolution of further resistance.

Microbial population dynamics often play out in fragmented habitats, yet our strategies for controlling microbes are largely modelled on well-mixed populations and often fail. Our study reveals some of the mechanisms that may underly such failure, and points to a need for deeper understanding of how microbial interactions and antibiotic therapy act in realistic contexts.

## Supporting information

Supplemental numerical methods, theory and results

## Acknowledgments

The authors thank Helen Alexander, Naama Brenner, Joel Ching Kuma Mbanghanih, Diana Fusco, Tatjana Malycheva, Daniel Taylor and Stefan Schuster for valuable discussions. The schematic illustrations in figures 2–5 were created using Biorender.com. NV and RJA were supported by the Deutsche Forschungsgemeinschaft (DFG) via the Excellence Cluster Balance of the Microverse (EXC 2051 - Project-ID 390713860) and via SFB 1127/2 ChemBioSys - 239748522 (Project C07). NV was also funded by the EPSRC Centre for Doctoral Training in Soft and Functional Interfaces (SOFI: EP/L015536/1). OP was funded by the EPSRC Centre for Doctoral Training in Soft Matter for Formulation and Industrial Innovation (SOFI^2^: EP/S023631/1). RJA was also funded by the European Research Council under Consolidator grant 682237 EVOSTRUC. For the purpose of open access, the author has applied a Creative Commons Attribution (CC BY) licence to any Author Accepted Manuscript version arising from this submission.

## Footnotes

1 For Poisson distributed initial numbers, the subpopulation density has mean *ρ* and variance *ρm*/*V*, hence the coefficient of variation 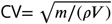

## References

Ackermann, M. (2015). A functional perspective on phenotypic heterogeneity in microorganisms. Nat. Rev. Microbiol., 13:497–508.

Alexander, H. K. and MacLean, R. C. (2020). Stochastic bacterial population dynamics restrict the establishment of antibiotic resistance from single cells. Proc. Natl. Acad. Sci. USA, 117:19455–19464.

Allen, R. J. and Waclaw, B. (2019). Bacterial growth: a statistical physicist’s guide. Rep. Prog. Phys., 82:016601.

Arnosti, C. (2011). Microbial extracellular enzymes and the marine carbon cycle. Annu. Rev. Mar. Sci., 3:401–425.

Artemova, T., Gerardin, Y., Dudley, C., Vega, N. M., and Gore, J. (2015). Isolated cell behavior drives the evolution of antibiotic resistance. Mol. Sys. Biol., 11:822.

Avraham, R., Haseley, N., Brown, D., Penaranda, C., Jijon, H. B., Trombetta, J. J., Satija, R., Shalek, A. K., Xavier, R. J., Regev, A., and Hung, D. T. (2015). Pathogen cell-to-cell variability drives heterogeneity in host immune responses. Cell, 162:1309–1321.

Balaban, N. Q., Merrin, J., Chait, R., Kowalik, L., and Leibler, S. (2004). Bacterial persistence as a phenotypic switch. Science, 305:1622–1625.

Barizien, A., Suryateja Jammalamadaka, M. S., Amselem, G., and Baroud, C. N. (2019). Growing from a few cells: combined effects of initial stochasticity and cell-to-cell variability. J. R. Soc. Interface, 16:20180935.

Batsch, M., Guex, I., Todorov, H., Heiman, C. M., Vacheron, J., Vorholt, J. A., Keel, C., and van der Meer, J. R. (2024). Fragmented micro-growth habitats present opportunities for alternative competitive outcomes. bioRxiv: 10.1101/2024.01.26.577336.

Bickel, S. and Or, D. (2020). Soil bacterial diversity mediated by microscale aqueous-phase processes across biomes. Nat. Commun., 11:116.

Bjarnsholt, T., Jensen, P., Moser, C., and Høiby, N. e. (2011). Biofilm infections. Springer, New York.

Blackburn, N., Fenchel, T., and Mitchell, J. G. (1998). Microscale nutrient patches in planktonic habitats shown by chemotactic bacteria. Science, 282:2254–2256.

Blume-Peytavi, U., Tosti, A., Whiting, D. A., and Trüeb, R. (2008). Hair growth and disorders. Springer-Verlag.

Boedicker, J. Q., Vincent, M. E., and Ismagilov, R. F. (2009). Microfluidic confinement of single cells of bacteria in small volumes initiates high-density behavior of quorum sensing and growth and reveals its variability. Angew. Chemie Int. Ed., 48:5908–5911.

Brook, I. (1989). Inoculum effect. Rev. Inf. Dis., 11:361–368.

Bush, K. and Bradford, P. A. (2016). β-lactams and β-lactamase inhibitors. Cold Spring Harb Perspect Med., 6:a025247.

CDC (2019). Antibiotic resistance threats in the United States. Technical report, U.S. Department of Health and Human Services.

Chuang, J. S., Rivoire, O., and Leibler, S. (2009). Simpson’s paradox in a synthetic microbial system. Science, 323:272–275.

Coates, J., Park, B. R., Le, D., Şimşek, E., Chaudry, W., and Kim, M. (2018). Antibiotic-induced population fluctuations and stochastic clearance of bacteria. eLife, 7:e32976.

Collins, D. J., Neild, A., DeMello, A., Liu, A.-Q., and Ai, Y. (2015). The Poisson distribution and beyond: methods for microfluidic droplet production and single cell encapsulation. Lab Chip, 15(17):3439–3459.

Connell, J. L., Kim, J., Shear, J. B., Bard, A. J., and Whiteley, M. (2014). Real-time monitoring of quorum sensing in 3D-printed bacterial aggregates using scanning electrochemical microscopy. Proc. Natl. Acad. Sci. USA, 111:18255–18260.

Conwill, A., Kuan, A. C., Damerla, R., Poret, A. J., Baker, J. S., Tripp, A. D., Alm, E. J., and Lieberman, T. D. (2022). Anatomy promotes neutral coexistence of strains in the human skin microbiome. Cell Host Microbe, 30:171–182.e7.

Dal Co, A., van Vliet, S., Kiviet, D. J., Schlegel, S., and Ackermann, M. (2020). Short-range interactions govern the dynamics and functions of microbial communities. Nature Ecol. Evol., 4:366–375.

Dann, L. M., Mitchell, J. G., Speck, P. G., Newton, K., Jeffries, T., and Paterson, J. (2014). Virio- and bacterioplankton microscale distributions at the sediment-water interface. PLoS ONE, 9:1–14.

Deris, J. B., Kim, M., Zhang, Z., Okano, H., Hermsen, R., Groisman, A., and Hwa, T. (2013). The innate growth bistability and fitness landscapes of antibiotic-resistant bacteria. Science, 342:1237435.

Dugatkin, L. A., Perlin, M., Lucas, J. S., and Atlas, R. (2005). Group-beneficial traits, frequency-dependent selection and genotypic diversity: an antibiotic resistance paradigm. Proc. R. Soc. B, 272:79–83.

Ebrahimi, A., Schwartzman, J., and Cordero, O. X. (2019). Cooperation and spatial self-organization determine rate and efficiency of particulate organic matter degradation in marine bacteria. Proc. Natl. Acad. Sci. USA, 116:23309–23316.

Edwards, J., Johnson, C., Santos-Medellin, C., Lurie, E., Kumar Podishetty, N., Bhatnagar, S., Eisen, J. A., and Sundaresan, V. (2015). Structure, variation and assembly of the root-associated microbiomes of rice. Proc. Natl. Acad. Sci. USA, 112:911–920.

Galloway, D. R. (1991). Pseudomonas aeruginosa elastase and elastolysis revisited: recent developments. Mol. Microbiol., 5:2315–2321.

Geyrhofer, L. and Brenner, N. (2020). Coexistence and cooperation in structured habitats. BMC Ecol., 20:14.

Geyrhofer, L., Ruelens, P., Farr, A. D., Pesce, D., de Visser, A. G. M., and Brenner, N. (2023). Minimal surviving inoculum in collective antibiotic resistance. mBio, 14:02456–22.

Gillespie, D. T. (1976). A general method for numerically simulating the stochastic time evolution of coupled chemical reactions. J. Comp. Phys., 22:403–434.

Gore, J., Youk, H., and van Oudenaarden, A. (2009). Snowdrift game dynamics and facultative cheating in yeast. Nature, 459:253–256.

Grinberg, M., Orevi, T., Steinberg, S., and Kashtan, N. (2019). Bacterial survival in microscopic surface wetness. Science, 8:e48508.

Hansen, R. H., Timm, A. C., Timm, A. N., Morrell-Falvey, J. L., Pelletier, D. A., Simpson, M. L., Doktycz, M. J., and Retterer, S. T. (2016). Stochastic assembly of bacteria in microwell arrays reveals the importance of confinement in community development. PLoS One, 11:e0155080.

Hermenau, R., Kugel, S., Komor, A. J., and Hertweck, C. (2020). Helper bacteria halt and disarm mushroom pathogens by linearizing structurally diverse cyclolipopeptides. Proc. Natl. Acad. Sci. USA, 117:23802—-23806.

Hsu, R. H., Clark, R. L., Tan, J. W., Ahn, J. C., Gupta, A., Romero, P. A., and Venturelli, O. S. (2019). Microbial interaction network inference in microfluidic droplets. Cell Systems, 9:229–242.

Khatri, B., Free, A., and Allen, R. J. (2012). Oscillating microbial dynamics driven by small populations, limited nutrient supply and high death rates. J. Theor. Biol., 314:120–129.

Kuzyakov, Y. and Blagodatskaya, E. (2015). Microbial hotspots and hot moments in soil: Concept & review. Soil Biol. Biochem., 83:184–199.

Mant, D., Orevi, T., and Kashtan, N. (2024). The impact of micro-habitat fragmentation on microbial populations growth dynamics. bioRxiv: 10.1101/2024.04.05.588087.

Meredith, H. R., Srimani, J. K., Lee, A. J., Lopatkin, A. J., and You, L. (2015). Collective antibiotic tolerance: mechanisms, dynamics and intervention. Nat. Chem. Biol., 11:182.

Mizrahi, S. P., Goyal, A., and Gore, J. (2022). Community interactions drive the evolution of antibiotic tolerance in bacteria. Proc. Natl. Acad. Sci. USA, 120(3).

Monier, J. and Lindow, S. E. (2004). Frequency, Size, and Localization of Bacterial Aggregates on Bean Leaf Surfaces. Appl. Environ. Microbiol., 70.

Ochs, M., Nyengaard, J. R., Jung, A., Knudsen, L., Voigt, M., Wahlers, T., Richter, J., and Gundersen, H. J. G. (2004). The number of alveoli in the human lung. Am. J. Respir. Crit. Care Med., 169:120–124.

Ortega, F. E., Koslover, E. F., and Theriot, J. A. (2019). Listeria monocytogenes cell-to-cell spread in epithelia is heterogeneous and dominated by rare pioneer bacteria. eLife, 8:e40032.

Park, J., Kerner, A., Burns, M. A., and Lin, X. N. (2011). Microdroplet-enabled highly parallel co-cultivation of microbial communities. PLoS ONE, 6:e17019.

Peña, J. and Nöldeke, G. (2018). Group size effects in social evolution. J. Theor. Biol., 457:211–220.

Ray, K., Marteyn, B., Sansonetti, P., and Tang, C. M. (2009). Life on the inside: the intracellular lifestyle of cytosolic bacteria. Nat. Rev. Microbiol., 7:333–340.

Raynaud, X. and Nunan, N. (2014). Spatial ecology of bacteria at the microscale in soil. PLoS ONE, 9:e87217.

Scheler, O., Makuch, K., Debski, P. R., Horka, M., Ruszczak, A., Pacocha, N., Sozański, K., Smolander, O., Postek, W., and Garstecki, P. (2020). Droplet-based digital antibiotic screen reveals single-cell clinal heteroresistance in an isogenic bacterial population. Sci. Rep., 10:3282.

Tacconelli, E. (2017). Global priority list of antibiotic-resistant bacteria to guide research, discovery, and devel-opment of new antibiotics. Technical report, World Health Organization.

Taylor, D., Verdon, N., Lomax, P., Allen, R. J., and Titmuss, S. (2022). Tracking the stochastic growth of bacterial populations in microfluidic droplets. Phys. Biol., 19:026003.

Teimouri, H. and Kolomeisky, A. B. (2019). Theoretical investigation of stochastic clearance of bacteria: first-passage analysis. J. R. Soc. Interface, 16:20180765.

Tekwa, E. W., Nguyen, D., Loreau, M., and Gonzalez, A. (2017). Defector clustering is linked to cooperation in a pathogenic bacterium. Proc. R. Soc. B, 284:20172001.

Thapa, S., Mishra, J., Arora, N., Mishra, P., Li, H., ÓHair, J., Bhatti, S., and Zhou, S. (2020). Microbial cellulolytic enzymes: diversity and biotechnology with reference to lignocellulosic biomass degradation. Rev. Environ. Sci. Biotechnol., 19:621–648.

Tooke, C. L., Hinchliffe, P., Bragginton, E. C., Colenso, C. K., Hirvonen, V. H. A., Takebayashi, Y.,, and Spencer, J. (2019). β-lactamases and β-lactamase inhibitors in the 21st century. J. Mol. Biol., 431:3472–3500.

Vega, N. M. and Gore, J. (2014). Collective antibiotic resistance: mechanisms and implications. Curr. Opin. Microbiol., 21:28.

Welch, J. L. M., Hasegawa, Y., McNulty, N. P., Gordon, J. I., and Borisy, G. G. (2017). Spatial organization of a model 15-member human gut microbiota established in gnotobiotic mice. Proc. Natl. Acad. Sci. USA, 114:9105–9114.

West, S. A., Griffin, A. S., Gardner, A., and Diggle, S. P. (2006). Social evolution theory for microorganisms. Nat. Rev. Microbiol., 4:597–607.

Wu, F., Ha, Y., Weiss, A., Wang, M., Letourneau, J., Wang, S., Luo, N., Huang, S., David, L. A., and You, L. (2022). Modulation of microbial community dynamics by spatial partitioning. Nature Chem. Biol., 18:394–402.

Yurtsev, E. A., Chao, H. X., Datta, M. S., Artemova, T., and Gore, J. (2013). Bacterial cheating drives the population dynamics of cooperative antibiotic resistance plasmids. Mol. Syst. Biol., 9:683.

Zhang, S., Mukherji, R., Chowdhury, S., Reimer, S., and Stallforth, P. (2021). Lipopeptide-mediated bacterial interaction enables cooperative predator defence. Proc. Natl. Acad. Sci. USA, 118:e2013759118.

Zhao, X., Ruelens, P., Farr, A. D., de Visser, A. G. M., and Baraban, L. (2023). Population dynamics of cross-protection against β-lactam antibiotics in droplet microreactors. Front. Microbiol., 14:1294790.

