## Supplemental numerical methods, theory and results for "Habitat fragmentation enhances microbial collective defence"

### Supplementary numerical simulation methods

To generate Figs. 3 and 4 in the main text, 1000 replicate simulations were used to calculate the survival fraction for each data point. Survival was defined by whether any subvolume had a surviving population determined after 300 minutes of simulated time.

#### Simulation of bacterial dynamics

In our numerical simulations, we model exponential bacterial growth or death via a simple timestepping procedure: in each timestep, $dt$, the population size $N(t)$ is decreased by $dN = -N \times \gamma \times dt$, where $\gamma$ is the death rate, or increased by $dN = N \times \mu \times dt$, where $\mu$ is the growth rate. In all simulations with deterministic population dynamics (except for Fig. S3b), the timestep $dt = 1$ min. To account for the discreteness of individual bacteria, the population size is set to zero if the number of bacteria $N(t)$ becomes less than 1.

#### Simulation of antibiotic dynamics

In our model, the degradation of antibiotic follows Michaelis-Menten kinetics:

$$\frac{da}{dt} = -\frac{r_{\mathrm{max}} a(t)}{a(t) + K_{\mathrm{M}}} R \,, \tag{S1}$$

where $R$ is a constant equal to the assumed enzyme concentration $bN/V$ ($b$ being the number of enzyme molecules per cell). To account accurately for the antibiotic degradation during one timestep $dt$ of our simulation, we integrate Eq. S1 over a timestep, i.e. from time $t$ to $t + dt$, as follows. We first separate the variables to reach

$$\int_{a(t)}^{a(t+dt)} \left(1 + \frac{K_{\mathrm{M}}}{a(t)}\right) da = \int_{t}^{t+dt} r_{\mathrm{max}} R \, dt \,. \tag{S2}$$

Performing the integral on both sides we obtain

$$\frac{a(t+dt)}{K_{\mathrm{M}}} + \log\left(a(t+dt)\right) = \frac{a(t)}{K_{\mathrm{M}}} + \log\left(a(t)\right) - \frac{r_{\mathrm{max}} R}{K_{\mathrm{M}}} dt \,. \tag{S3}$$

By exponentiating we reach

$$\frac{a(t+dt)}{K_{\mathrm{M}}} e^{\frac{a(t+dt)}{K_{\mathrm{M}}}} = \frac{a(t)}{K_{\mathrm{M}}} e^{\frac{a(t)}{K_{\mathrm{M}}}} e^{-\frac{r_{\mathrm{max}} R \tau}{K_{\mathrm{M}}}} \,. \tag{S4}$$

The solution for $a(t+dt)$ can be expressed in terms of the Lambert W function (***Abramowitz and Stegun, 1965***):

$$a(t+dt) = K_{\mathrm{M}} \times W\left(\frac{a(t)}{K_{\mathrm{M}}} e^{\frac{a(t)}{K_{\mathrm{M}}}} \times e^{-\left(\frac{r_{\mathrm{max}} b N(t)}{K_{\mathrm{M}} V}\right) dt}\right) , \tag{S5}$$

where we have substituted back the expression for $R$. Since we are dealing with positive real numbers, we choose the principal branch (the principal branch $W(z)$ is real when $z < -1/e$).

#### Stochastic allocation of bacteria to subvolumes

To mimic stochastic allocation of bacteria amongst the subvolumes, the initial number of bacteria for a given simulation run is selected from a Poisson distribution using the function poisson.rvs in the python package scipy.stats. The parameter of this Poisson distribution corresponds to the average number of bacteria per subvolume, which is calculated based on the subvolume size $v$ and the assumed density of bacteria in the bulk $\rho$ (in this work, $\rho = 5 \times 10^{-7}$ cells/ml).

#### Enzymatic defence: parameters

The parameters used in our simulations are listed in Table S1.

| Parameter | Value | Unit | Source |
|---|---|---|---|
| $K_M$ | 6.7 | $\mu$g /ml | ***Yurtsev et al.*** (***2013***) |
| $br_{max}$ | $3.5 \times 10^{-8}$ | $\mu$g/cell/min | ***Yurtsev et al.*** (***2013***) |
| $a_{\text{th}}$ | 1 | $\mu$g/ml | ***Yurtsev et al.*** (***2013***) |
| $V$ | $1 \times 10^{-4}$ | ml | ***Taylor et al.*** (***2022***) |
| $dt$ | 1 | min | determined by simulation trials |
| $\gamma$ | 0.045 | /min | ***Yurtsev et al.*** (***2013***) |
| $\mu$ | 0.01 | /min | ***Verdon*** (***2023***) |

**Table S1. Default parameters used in our simulations.** The unfragmented volume $V$ is chosen to be 1000 times larger than the volume of a single droplet in a typical microfluidic experiment, e.g. that of ***Taylor et al.*** (***2022***).

#### Enzymatic defence: stochastic simulations

To model the effects of intrinsic stochasticity in microbial dynamics for the enyzmatic defence model (i.e. for the simulations shown in Fig. S2b), we used the Gillespie algorithm (***Gillespie, 1976***). In these simulations, microbial birth and death events are assumed to occur as Markov processes with rate $\mu$ and $\gamma$ respectively, with only death events occurring if the antibiotic concentration is above the threshold, and only birth events occurring if the antibiotic concentration is below the threshold. The antibiotic concentration $a(t)$ was modelled deterministically (since the absolute number of antibiotic molecules is typically much larger than the number of microbes), using the analytical solution for the change in $a(t)$ from one timestep to the next (Eq. S5).

### Supplementary theoretical derivations

#### Assumption of mixing within a subvolume

A key assumption of our model is that the system is well-mixed within each subvolume, so that we do not need to consider the spatial arrangement of bacteria and/or chemicals within the subvolume. To justify this assumption, we consider here the typical time required for an antibiotic molecule to diffuse across a single subvolume.

The diffusion constant $D$ of an antibiotic molecule can be estimated using the Stokes-Einstein relation: $D = k_B T/(6\pi\eta r)$, where $r$ is the molecular radius and $\eta$ is the viscosity of the media. We estimate the radius of an ampicillin molecule, $r$, as 10Å, we assume a temperature, $T$, of 37°C (310 K) and we use the viscosity, $\eta$, of water at this temperature ($7 \times 10^{-4}$ Pa s). This results in a diffusion constant $D = 3.2 \times 10^{-10}$ m$^2$s$^{-1}$. For diffusive motion the mean square displacement $msd$ scales linearly with time $\tau$: msd$(\tau) = 6D\tau$. In our simulations, the subvolume $v$ for degree of habitat fragmentation $m = 1000$ is $v = 100$ pl $= 10^{-10}$ l, corresponding to a sphere of radius approximately 35 $\mu$m (of the order of the size of a cell). The time $\tau$ that is required for an ampicillin molecule to diffuse this distance is then $(35{\times}10^{-6})^2/(6{\times}3.2{\times}10^{-10}) = 0.6$s. For the largest volume that we simulate ($m = 1$), we have $v = 10^{-7}$ l, corresponding to a sphere of radius $\sim 300\mu$m, and hence $\tau \approx 60$s. Since the timescales of bacterial growth and death are $1/\mu = 100$ min and $1/\gamma = 22$ min respectively, we conclude that the timescale for antibiotic diffusion is much shorter that that of the bacterial growth or death dynamics. We also note that our estimate for the antibiotic mixing timescale is conservative, since bacterial motility may also contribute to active mixing within a subvolume.

### Enzymatic defence: solution of the deterministic collective defence model and expression for $\rho^*$

We first recap the defining equations for our collective enzymatic defence model. The dynamics of the bacterial population obey

$$\dot{N}(t) = N(t)\left[\mu\theta(a_{\rm th} - a) - \gamma\theta(a - a_{\rm th})\right] \, , \tag{S6}$$

where $\theta(x)$ is the Heaviside step function, $\mu$ is the growth rate for low antibiotic $a(t) < a_{\rm th}$, and $\gamma$ is the death rate for high antibiotic $a(t) > a_{\rm th}$. The dynamics of the antibiotic concentration $a(t)$ obey Eq. S1, in which it is assumed that each enzyme degrades antibiotic according to Michaelis-Menten kinetics, i.e. with the rate per enzyme molecule being $r_{\rm max}a(t)/(a(t)+K_{\rm M})$. Here, $r_{\rm max}$ and $K_{\rm M}$ are the Michaelis-Menten parameters: respectively, the maximal rate of degradation per enzyme, and the antibiotic concentration at which the rate of degradation per enzyme is half-maximal. As mentioned in the main text, if enzyme molecules are not fully secreted but instead remain wholly or partially inside the cell, the parameter $r_{\rm max}$ implicitly includes a factor accounting for transport of antibiotic across the cell boundary. We assume that the initial antibiotic concentration, $a_{\rm init}$, is high, $a_{\rm init} > a_{\rm th}$, so that the population is initially killed. Similar models have been proposed in previous studies (***Yurtsev et al., 2013***; ***Mizrahi et al., 2022***; ***Geyrhofer et al., 2023***). In particular the model of ***Geyrhofer et al.*** (***2023***) uses a more detailed form of the antibiotic concentration-dependent growth and killing rates, but reaches similar conclusions regarding the two possible fates of the microbial population and the density-dependent conditions for survival.

Eqs. S6 and S1 can be solved to predict the fate of the bacterial population. If the initial antibiotic concentration is higher than $a_{\rm th}$, from Eqs. S6 we find that the population initially decreases exponentially as $N(t) = N_{\rm init}e^{-\gamma t}$. Substituting this into Eq. S1 and integrating shows that the dynamics of the antibiotic concentration obeys

$$\frac{F[a(t), a_{\rm init}]}{r_{\rm max}} = \frac{bN_{\rm init}}{\gamma V}\left[1 - e^{-\gamma t}\right] \, , \tag{S7}$$

where $F[a(t), a_{\rm init}] \equiv \left[(a_{\rm init} - a(t)) + K_{\rm M}\ln\left(a_{\rm init}/a(t)\right)\right]$.

If the antibiotic concentration decreases below the threshold $a_{\rm th}$, the bacterial population will start to regrow and will ultimately survive. Denoting the time $t_{\rm th}$ such that $a(t_{\rm th}) = a_{\rm th}$ and using our result (S7) for the dynamics of the antibiotic concentration, we find that

$$t_{\rm th} = -\frac{1}{\gamma}\ln\left[1 - \frac{\gamma V}{bN_{\rm init}r_{\rm max}}F[a_{\rm th}, a_{\rm init}]\right] . \tag{S8}$$

We can then write the full expression for the dynamics of the microbial population:

$$N(t) = N_{\rm init}e^{-\gamma t} \qquad t \leq t_{\rm th} \, ; \tag{S9}$$

$$N(t) = N(t_{\rm th})e^{\mu(t-t_{\rm th})} \qquad t > t_{\rm th} \, , \tag{S10}$$

where

$$N(t_{\rm th}) = N_{\rm init} - \frac{\gamma V}{br_{\rm max}}F[a_{\rm th}, a_{\rm init}]. \tag{S11}$$

The full expression for the population size at long times, after regrowth, is then (from Eqs. S8, S10 and S11):

$$N(t) = N_{\rm init}e^{\mu t}\left[1 - \frac{\gamma V F[a_{\rm th}, a_{\rm init}]}{bN_{\rm init}r_{\rm max}}\right]^{\left(1+\frac{\mu}{\gamma}\right)} . \tag{S12}$$

Using Eq. S8, we can determine how the fate of the bacterial population depends on the parameters of the model. As the argument of the logarithm in Eq. S8 tends to zero, the time $t_{\rm th}$ at which the antibiotic concentration reaches $a_{\mathrm{th}}$ tends to infinity, implying that the antibiotic concentration never reaches the threshold, and hence that the population does not regrow, i.e. it is killed. For the population to survive and eventually regrow we require $t_{\mathrm{th}}$ to be finite. In other words, the condition for survival is $N_{\mathrm{init}}/V > \rho^*$, where the threshold initial population density $\rho^*$ is as given in the main text:

$$\rho^* = \frac{\gamma}{br_{\max}} \left[ (a_{\mathrm{init}} - a_{\mathrm{th}}) + K_{\mathrm{M}} \ln \left( \frac{a_{\mathrm{init}}}{a_{\mathrm{th}}} \right) \right] . \tag{S13}$$

Eq. S13 defines the phase boundary line between the survival and killing regions of the parameter space, as discussed in the main text and shown in Fig. 2c.

### Enzymatic defence: scaling of survival probability with degree of habitat fragmentation

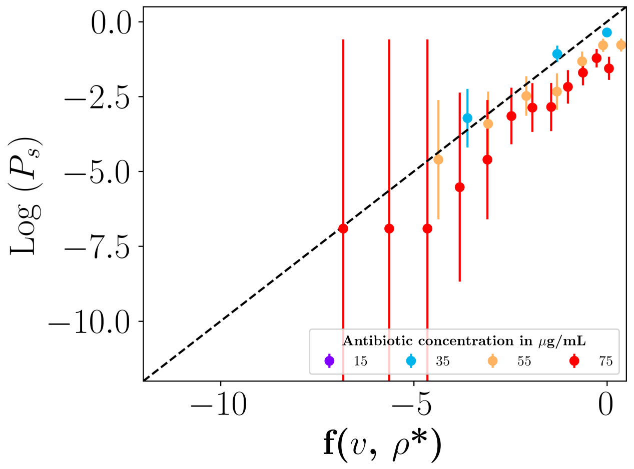


**Figure S1. Scaling of survival probability $P_s$.** The logarithm of the survival probability, $\log P_s$, obtained from our simulations using deterministic population dynamics with Poisson distributed initial population size (as in Fig. 4 of the main text), is plotted as a function of the right hand side of Eq. S17. We denote $f(v, \rho^*) \equiv \rho^* v\,(1 + \log(\rho/\rho^*) - \rho/\rho^*) + \log(V/v) - \frac{1}{2}\log(2\pi\rho^* v)$. The data points collapse onto a straight line with gradient unity (dashed line), as predicted by Eq. S17. The small offset of the simulation data from the theoretical line may arise from the approximations made in deriving Eq. S17.

In our model, microbes are allocated stochastically to subvolumes, so the subpopulation sizes are Poisson-distributed. Stochastic partitioning of antibiotic molecules is not considered, since the absolute number of antibiotic molecules at these concentrations is far higher than the number of microbes.

Assuming deterministic population dynamics, a subpopulation survives if its initial population density is higher than the threshold $\rho^*$. The initial number $N_{i,\mathrm{init}}$ is Poisson distributed with mean $\rho v = \rho V/m$, so the survival probability for a subpopulation is

$$p_s = e^{-\rho v} \sum_{j=N^*}^{\infty} \frac{(\rho v)^j}{j!} , \tag{S14}$$

where $N^* = \lceil \rho^* v \rceil$. We are interested in the case where the bulk population density $\rho$ is below the survival threshold: $\rho < \rho^*$ (i.e. the killing part of the phase diagram, Fig. 2c of the main text, below the phase boundary). In this case the largest contribution to the sum in Eq. S14 comes from the first term. We therefore approximate the sum by the first term: $p_s \approx e^{-\rho v}(\rho v)^{N^*}/N^*!$. Using Stirling's approximation $N^*! \approx \sqrt{2\pi N^*}(N^*/e)^{N^*}$ we then obtain $p_s \approx \frac{e^{-\rho v}}{\sqrt{2\pi N^*}} \left(\frac{\rho v e}{N^*}\right)^{N^*}$, and setting $N^* \approx \rho^* v$ this becomes

$$p_s \approx \frac{e^{-\rho v}}{\sqrt{2\pi\rho^* v}} \left(\frac{\rho e}{\rho^*}\right)^{\rho^* v} = \frac{e^{\rho^* v(1+\log(\rho/\rho^*) - (\rho/\rho^*))}}{\sqrt{2\pi\rho^* v}} . \tag{S15}$$

The survival probability $P_s$ for the entire population is given by $P_s = 1 - (1 - p_s)^m$. For small $p_s$, this can be approximated as $P_s \approx 1 - (1 - mp_s) = mp_s$. Using $m = V/v$, we then obtain

$$P_s \approx \frac{V}{v}\left(\frac{e^{\rho^* v(1+\log(\rho/\rho^*)-\rho/\rho^*)}}{\sqrt{2\pi\rho^* v}}\right) \tag{S16}$$

(where $\log$ denotes the natural logarithm). Therefore we expect $\log(P_s)$ to scale with $v$ as

$$\log P_s \approx \rho^* v\left(1 + \log\frac{\rho}{\rho^*} - \frac{\rho}{\rho^*}\right) + \log\left(\frac{V}{v}\right) - \frac{1}{2}\log(2\pi\rho^* v)\,. \tag{S17}$$

For small $v$ we expect a linear relationship between $\log P_s$ and $v$. Fig. S1 shows that, indeed, in our simulations and theory, the data collapses when $\log P_s$ is plotted versus the right-hand-side of Eq. S17.

### Supplementary results

#### Enzymatic defence: habitat fragmentation increases total population size under lethal conditions

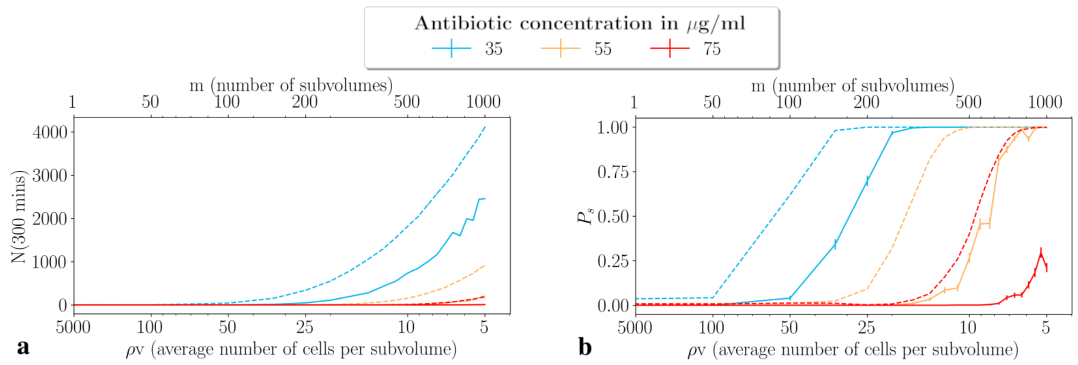


**Figure S2. Total population size increases with degree of habitat fragmentation under lethal conditions, and demographic stochasticity can also lead to habitat-fragmentation rescue**. (a) Simulation results for the total population size $N_{\mathrm{tot}}$ at time $t = 300$ min, for different degrees of habitat fragmentation. Results are shown for several concentrations of antibiotic (indicated by coloured lines; see legend). The solid lines show simulations with Poisson partitioning and deterministic population dynamics; the dashed lines show simulations with Poisson partitioning and stochastic population dynamics. (b) Corresponding results for the population survival probability $P_s$. All simulations are averaged over 1000 replicate runs.

For a microbial population engaged in collective enzymatic defence, we show in the main text that habitat fragmentation allows survival under lethal conditions for which a well-mixed, non-fragmented population would be killed ($\rho < \rho^*$). It is also interesting to predict the dynamics of the total population size $N_{\mathrm{tot}}(t)$ for such populations, at long times.

On short timescales, we expect to see killing in every subpopulation – i.e. the population size $N_i(t)$ in subvolume $i$ decreases exponentially at rate $\gamma$ for all subvolumes $i$. On longer timescales, however, different subpopulations may have different fates. In some subvolumes the initial population density will be below the survival threshold, $N_{i,\mathrm{init}}/v < \rho^*$. These subpopulations will die, hence they will not contribute to the total population size $N_{\mathrm{tot}}$ at long times. To compute $N_{\mathrm{tot}}$ at long times, we therefore only need to consider the long-time dynamics $N_i(t)$ of those subpopulations which have initial density above the survival threshold: $N_{i,\mathrm{init}}/v > \rho^*$. By analogy with Eq. S12 (or using Eqs. S8, S10 and S11, this is

$$N_i(t) \quad = \quad N_{i,\mathrm{init}} e^{\mu t}\left[1 - \frac{\gamma v F[a_{\mathrm{th}}, a_{\mathrm{init}}]}{b N_{i,\mathrm{init}} r_{\mathrm{max}}}\right]^{(1+\mu/\gamma)} . \tag{S18}$$

Eq. S18 shows that the growth of each surviving subpopulation is exponential with rate $\mu$, but with a prefactor that depends on the initial population size $N_{i,\text{init}}$. This arises because the time $t_{\text{th}}$ at which regrowth of subpopulation $i$ starts, and the corresponding population size $N_i(t_{\text{th}})$ from which regrowth starts, depend on $N_{i,\text{init}}$.

The total population size $N_{\text{tot}}(t)$ can be found by summing Eq. S18 over all initial population sizes larger than the survival threshold $\rho^* v$, weighted by their Poisson probability, and multiplying by the total number of subvolumes $m$:

$$N_{\text{tot}}(t) = e^{\mu t} \times e^{-\rho v} \times m \sum_{j=N^*}^{\infty} \frac{(\rho v)^j}{j!} \times j \left[1 - \frac{\gamma v F[a_{\text{th}}, a_{\text{init}}]}{j b r_{\text{max}}}\right]^{(1+\mu/\gamma)}, \tag{S19}$$

where $N^* = \lceil \rho^* v \rceil$.

We are interested in the effect of habitat fragmentation on $N_{\text{tot}}(t)$. Fig. S2 shows the total population size $N_{\text{tot}}$, measured at a fixed time $t = 300$ min, for different degrees of fragmentation $m$ (we note that $m$ also enters Eq. S19 through the subvolume $v = V/m$ and through the lower bound $N^*$ in the sum), for several antibiotic concentrations, and for both deterministic and stochastic population dynamics. As expected, habitat fragmentation increases the total population size. This is because subvolumes whose initial density is further above the survival threshold $\rho^*$ start regrowth earlier and contribute disproportionately to the final population size.

### Limit of the model for extreme habitat fragmentation

As discussed in the main text, our model predicts that for extreme habitat fragmentation, where occupied subvolumes become so small that the local density is above $\rho^*$ even for a single microbe, the subpopulation survival probability $p_s$ approaches 1. In this limit, the model also suggests that the population size increases exponentially with the same dynamics as a bulk, unfragmented population.

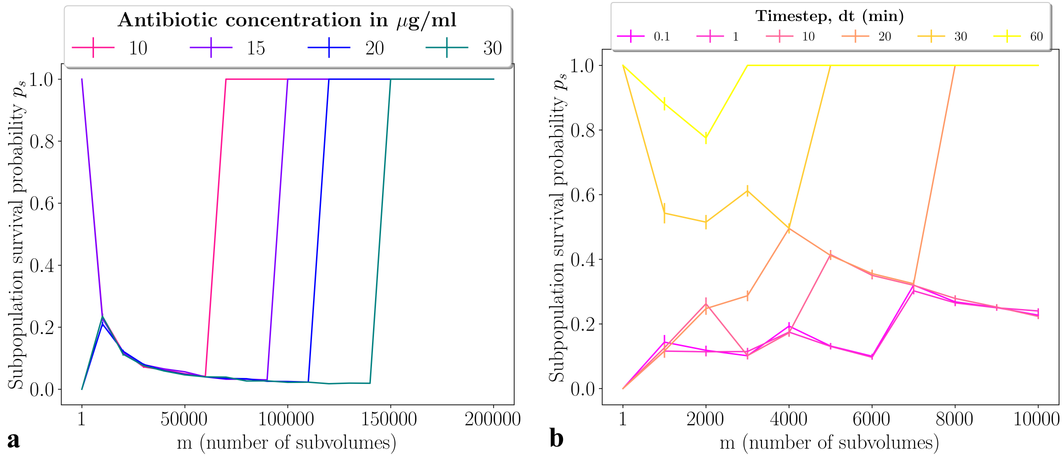


**Figure S3. Model predictions for extreme habitat fragmentation (large m).** (a): Subpopulation survival probability $p_s$ as a function of degree of habitat fragmentation $m$ for extreme fragmentation, for several initial antibiotic concentrations (indicated by colours). This represents an extension of Fig. 5 of the main text to more extreme fragmentation. For extreme fragmentation the subvolumes become so small that bacterial density in each subvolume is above the threshold $\rho^*$, so $p_s$ tends to 1. (b): Timestep dependence of simulation results for extreme fragmentation. The subpopulation survival probability $p_s$ is shown for different values of the simulation timestep $dt$ (indicated by colours). Here, $a_{\text{init}} = 30\mu$g/ml and $\rho = 5 \times 10^7$ cells/ml; hence the threshold density $\rho^*$ =$6.66 \times 10^7$ cells/ml. The theory predicts that $p_s = 1$ for values of $m$ larger than $6659$ (corresponding to $v = 1/\rho^*$).

To see this mathematically, we note that in the limit of extreme habitat fragmentation, where there is only 1 or 0 microbes per subvolume, the sum in Eq. S19 has only one non-zero contribution, for $j = 1$. Eq. S19 then reduces to

$$\begin{aligned} N_{\mathrm{tot}}(t) &= e^{\mu t} \times e^{-\rho v} \times \rho v m \left[1 - \frac{\gamma v F[a_{\mathrm{th}}, a_{\mathrm{init}}]}{b r_{\max}}\right]^{(1+\mu/\gamma)} \\ &= e^{\mu t} \times e^{-\frac{N_{\mathrm{init}}}{m}} \times N_{\mathrm{init}} \left[1 - \frac{\gamma V F[a_{\mathrm{th}}, a_{\mathrm{init}}]}{m b r_{\max}}\right]^{(1+\mu/\gamma)} \\ &\to N_{\mathrm{init}} e^{\mu t}\,, \end{aligned} \tag{S20}$$

where we have used the fact that the initial density $\rho = N_{\mathrm{init}}/V$ and taken the limit of large $m$, i.e. small $1/m$. Thus, in the limit of very high degree of fragmentation, the dynamics reduces to uninhibited, antibiotic-free, exponential growth. This is because every microbe finds itself in a tiny subvolume containing few antibiotic molecules, which are degraded rapidly (within each occupied subvolume, $t_{\mathrm{th}} \to 0$).

This model prediction must, however, be viewed with caution. Our model treats the microbial population density as a continuous variable, but this assumption falls down when we approach the limit of a single microbe. In our numerical simulations, we account for the discreteness of individual microbes by setting the population size $N(t)$ to zero when $N(t) < 0$. From a numerical point of view, this results in an apparent timestep dependence of the subpopulation survival probability for extreme fragmentation (Fig. S3). This happens because during one timestep $dt$ the variable $N(t)$ may temporarily drop below 1. In other words, the outcome depends on the rate at which a single bacterium degrades antibiotic relative to the rate at which it is killed. Making quantitative predictions in this regime would require a more detailed understanding of the physiological factors that are at play when a single $\beta$-lactamase producing bacterium experiences a concentration of antibiotic that is above the scMIC.

### Enzymatic defence: stochastic birth and death dynamics favours survival and growth under lethal conditions

In the main text, we focus on the Poisson distribution of microbes among subvolumes as a source of stochasticity, leading to survival of enzyme-producers under lethal conditions. However, microbial birth and death dynamics are also intrinsically stochastic. This demographic stochasticity can also favour survival and regrowth under conditions where a well-mixed population would be killed ($\rho < \rho^*$). This is shown in Fig. S2, where we compare results for deterministic population dynamics those obtained with stochastic population dynamics, simulated with the Gillepsie algorithm ((***Gillespie, 1976***); see simulation methods section above). Fig. S2 shows that the addition of demographic stochasticity in our simulations enhances both the survival probability and the population size.

To understand the relative importance of the two source of stochasticity (initial population partitioning vs demographic stochasticity), we next performed a series of simulations, investigating the 4 scenarios shown in Fig. S4: deterministic partitioning and population dynamics (i.e. all subpopulations have the same initial population size and grow/die deterministically), deterministic partitioning and stochastic dynamics, stochastic partitioning and deterministic population dynamics, and both partitioning and population dynamics simulated stochastically. These simulations are performed for a parameter set ($\rho = 5 \times 10^7$ cells/ml, $a_{\mathrm{init}} = 35\mu$g/ml, hence $\rho^* = 7.4 \times 10^7$ cells/ml) for which a bulk population would be killed; hence the population survival probability $P_s = 0$ for the fully deterministic simulations. Stochasticity in either population dynamics or partitioning leads to survival for degree of habitat fragmentation $m \gtrsim 200$, while full stochasticity (in both population dynamics and partitioning) leads to population survival at a lower degree of habitat fragmentation ($m \gtrsim 100$). Therefore, for this parameter set, the contributions of the two sources of stochasticity are approximately equal in magnitude and are additive.

### Collective resource foraging model

In our collective resource foraging model, discussed in the main text, nutrients are released by enzyme-mediated degradation of an environmental substrate (e.g. a biopolymer). The microbial

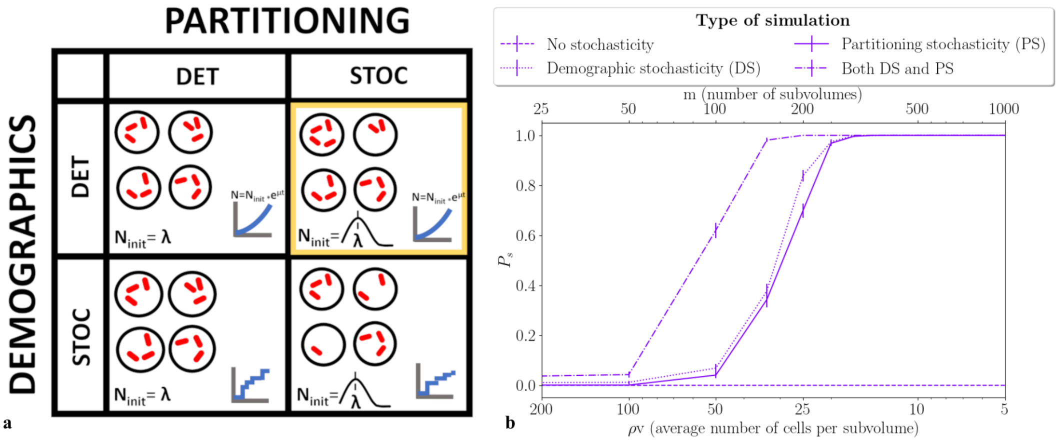


**Figure S4. Importance of demographic vs partitioning noise.** (a) Schematic illustration of the 4 types of simulations performed: deterministic partitioning with deterministic population dynamics, deterministic partitioning with stochastic dynamics, stochastic partitioning with deterministic population dynamics (highlighted in yellow: as used in the simulations for the main text), and stochastic partitioning with stochastic population dynamics. (b) Simulations results for the population survival probability $P_s$, plotted as a function of the degree of habitat fragmentation, for the 4 simulation types, for initial antibiotic concentration 35 $\mu$g/ml. Each data point represents the mean survival fraction of 1000 simulated replicates.

---

population cannot grow until a threshold concentration of nutrient has been released; this corresponds to a threshold concentration of the substrate having been degraded. Once this threshold is passed, we suppose that the population grows exponentially at a fixed rate. This model is simplistic: for example, it does not account for nutrient consumption by the growing microbes, so that it can predict only very early time growth dynamics. However, it does capture the essential feature of collective nutrient release from the environment.

The defining equations for the microbial population size $N(t)$ and the substrate (e.g. biopolymer) that is being degraded, $z(t)$, are

$$\dot{N}(t) = \mu N(t)\theta(z_{\text{th}} - z(t)), \tag{S21}$$

and

$$\dot{z}(t) = \frac{bN(t)}{V}\left(\frac{r_{\text{max}} z(t)}{z(t) + K_{\text{M}}}\right), \tag{S22}$$

where $\theta(x)$ denotes the Heaviside step function, $V$ is the total volume, $\mu$ is the microbial growth rate, $b$ is the number of substrate-degrading enzymes per microbe and $r_{\text{max}}$ and $K_{\text{M}}$ are the Michaelis-Menten parameters for enzymatic degradation of the substrate. The parameter values used in our simulations are listed in the caption of Fig. S5.

Fig. S5 shows simulation results for the dynamics of the normalised total population size $N_{\text{tot}}(t)/N_{\text{init}}$ in this model, for different degrees of habitat fragmentation $m$. The population initially does not grow, since not enough substrate has been degraded. After a lag phase, the threshold nutrient concentration is reached and exponential growth starts. Comparing results for different degrees of habitat fragmentation $m$ (denoted by colour in Fig. S5) we see that fragmentation leads to an earlier exit from the lag phase. This happens because subpopulations with an initially higher density degrade the substrate faster. Therefore habitat fragmentation provides an early-time growth advantage. On longer timescales, however, the population grows exponentially at the same rate ($\mu$) regardless of its degree of fragmentation. This is because all subpopulations eventually reach the threshold and start to grow. Once all subpopulations are growing, the total population size includes contributions both from initially more dense subpopulations (which started growing earlier) and initially less dense subpopulations (which started growing later). In the limit that the differences between subpopulations are small (i.e. the exponential difference in their growth dynamics can be

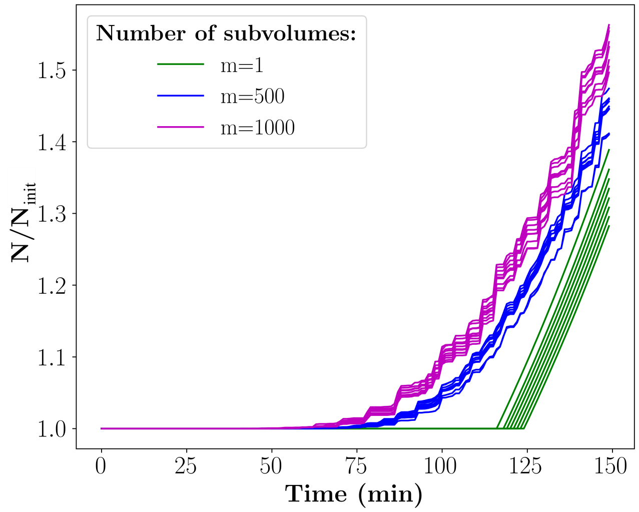


**Figure S5. Collective resource foraging model.** Simulation results for the collective resource foraging model defined by Eqs. S21 and S22, with Poisson distributed initial cell numbers. The total population size $N_{\mathrm{tot}}(t)$ is plotted, normalised by the initial total population size $N_{\mathrm{init}}$. The parameters are $z_{\mathrm{init}}$ =100 $\mu$g/ml, $z_{\mathrm{th}}$ =50 $\mu$g/ml, $K_{\mathrm{M}}$ =15 $\mu$g/ml, $br_{\mathrm{max}}$=$1 \times 10^{-8}$$\mu$g /cell/min and $\mu$ =0.01 /min. Results are shown for degree of habitat fragmentation $m = 1$ (mean initial population size per subvolume $\overline{N_{\mathrm{i,init}}} = 5000$ cells, $m = 500$ ($\overline{N_{\mathrm{i,init}}} = 10$) and $m = 1000$ ($\overline{N_{\mathrm{i,init}}} = 5$). In each case, 10 replicate simulations were performed for the entire population of $m$ subvolumes; each line in the plot shows one of the replicate simulations. The lag time for the $m = 1$ simulations shows some variability because the initial bacterial number is drawn from a Poisson distribution.

approximated by a linear function), these contributions tend to balance each other. In contrast, if the initial variance in subpopulation size is large, the contributions of initially more and less dense subvolumes do not balance in the long time limit, and there will be a lasting effect of habitat fragmentation on total population growth. This, however, a quantitative rather than a qualitative effect on the ecological outcome.

### References

Abramowitz, M. and Stegun, I. A. (1965). *Handbook of Mathematical Functions*. Dover.

Geyrhofer, L., Ruelens, P., Farr, A. D., Pesce, D., de Visser, A. G. M., and Brenner, N. (2023). Minimal surviving inoculum in collective antibiotic resistance. *mBio*, 14:02456–22.

Gillespie, D. T. (1976). A general method for numerically simulating the stochastic time evolution of coupled chemical reactions. *J. Comp. Phys.*, 22:403–434.

Mizrahi, S. P., Goyal, A., and Gore, J. (2022). Community interactions drive the evolution of antibiotic tolerance in bacteria. *Proc. Natl. Acad. Sci. USA*, 120(3).

Taylor, D., Verdon, N., Lomax, P., Allen, R. J., and Titmuss, S. (2022). Tracking the stochastic growth of bacterial populations in microfluidic droplets. *Phys. Biol.*, 19:026003.

Verdon, N. (2023). *Heterogeneous growth and death of small bacterial populations in microfuidic droplets*. PhD thesis, The University of Edinburgh.

Yurtsev, E. A., Chao, H. X., Datta, M. S., Artemova, T., and Gore, J. (2013). Bacterial cheating drives the population dynamics of cooperative antibiotic resistance plasmids. *Mol. Syst. Biol.*, 9:683.
